# Convergence and divergence in anti-predator displays: A novel approach to quantitative behavioural comparison in snakes

**DOI:** 10.1101/849703

**Authors:** Alison R. Davis Rabosky, Talia Y. Moore, Ciara M. Sánchez-Paredes, Erin P. Westeen, Joanna G. Larson, Briana A. Sealey, Bailey A. Balinski

## Abstract

Animals in nature use diverse strategies to evade or deter their predators, including many vivid behavioural displays only qualitatively described from field encounters with natural predators or humans. Within venomous snake mimicry, stereotyped anti-predator displays are suggested to be a critical component of the warning signal given by toxic models and thus under strong selection for independent convergence in mimetic species. However, no studies have systematically quantified variation in snake anti-predator displays across taxonomically broad clades to test how these behaviours evolve across species within a phylogenetic comparative methods framework. Here we describe a new, high-throughput approach for collecting and scoring snake anti-predator displays in the field that demonstrates both low observer bias and infinite extension across any species. Then, we show our method’s utility in quantitatively comparing the behaviour of 20 highly-divergent snake species from the Amazonian lowlands of Peru. We found that a simple experimental setup varying simulated predator cues was very successful in eliciting anti-predator displays across species and that high-speed videography captured a greater diversity of behavioural responses than described in the literature. We also found that although different display components evolve at different rates with complicated patterns of covariance, there is clear evidence of evolutionary convergence in anti-predator displays among distantly related elapid coral snakes and their colubrid mimics. We conclude that our approach provides new opportunity for analyses of snake behaviour, kinematics, and the evolution of anti-predator signals more generally, especially macroevolutionary analyses across clades with similarly intractable behavioural diversity.

## Introduction

As time-calibrated phylogenies across large taxonomic clades become increasingly available, behavioural ecologists have great opportunity to test macroevolutionary hypotheses about how and why behavioural traits evolve over time. However, models of trait evolution at this scale require the difficult task of 1) generating behavioural data for all of the taxa represented in the phylogeny, and 2) designing a quantification regime for traits that can accommodate the vast diversity, hierarchy, or non-equivalency of behavioural states that may exist across highly divergent taxa (Jablonski, 2017; O’Meara, 2012; Wainwright, 2007). Because interactions between predator and prey are fundamental drivers of many ecological and evolutionary dynamics in nature, anti-predator displays - in which animals use spectacular acoustic, chemical, and visual signals to avoid detection or consumption - have emerged as a prime target for testing theory about signal evolution more generally (Ruxton, Allen, Sherratt, & Speed, 2018). In particular, systems with large-scale diversity in both cryptic and warning signals provide a powerful opportunity to test hypotheses about the factors that promote the origin and maintenance of anti-predator traits that are convergent across phylogenetic lineages (Davis Rabosky et al., 2016).

Snakes are a compelling example of a clade with great potential for macroevolutionary analyses of behavioural traits, but that are substantially limited by the amount of data available (Greene, 1988). Most knowledge about snake anti-predator behaviour comes from a limited number of species, especially from North America (Arnold & Bennett, 1984; Brodie III, 1992; Greene, 1988; Herzog & Schwartz, 1990). Even in well-studied species, the extent to which defensive displays are affected by ambient temperature, lighting conditions, pheromones, or other stimuli remain a target of research (Brodie III & Russell, 1999; Ford, 1995; Schieffelin & de Queiroz, 1991; Shine, Olsson, Lemaster, Moore, & Mason, 2000). Broad surveys of anti-predator behaviour have been particularly difficult due to the relative rarity and/or low detectability in snakes, especially in the high richness areas of the tropical latitudes where we simply have no data for most species (Martins, 1996). The extreme humidity, rainfall, and heat conditions of the tropics have also posed technical challenges for high speed video equipment necessary to record rapid snake behaviours, limiting most work to laboratory tests at universities (Ford, 1995). However, recent developments in affordable, yet durable, solar power and video technology mean that comparative biologists looking to maximise data collection on anti-predator behaviour across a large portion of the snake phylogeny can now go anywhere in the world under nearly any field conditions and create a permanent video record of a snake’s response to a simulated or real predator cue in nature.

### The role of behaviour in Batesian mimicry

Snakes play an important role in animal behaviour because of their contribution to understanding Batesian mimicry, in which toxic or noxious species are imitated by harmless species to deter predation (Bates, 1862; Wallace, 1867; Wickler, 1968). In the Western Hemisphere, the bright red-and-black coloration of highly venomous coral snakes (genus *Micrurus*) in the family Elapidae is mimicked by at least 150 species of distantly-related snakes (Campbell & Lamar, 2004; Davis Rabosky et al., 2016; Savage & Slowinski, 1992) in the families Colubridae (*sensu stricto* following Pyron, Burbrink, & Wiens, 2013) and Aniliidae. Analysis of coloration is a well-understood metric for assessing mimetic potential, as nearly fifty years of experimental research has demonstrated that predators innately avoid “coral snake-like” colour patterns on clay replicas and that this avoidance provides a selective advantage over other colour patterns (Akcali & Pfennig, 2014; Brodie III, 1993; Brodie III & Janzen, 1995; Pfennig, Harcombe, & Pfennig, 2001; Smith, 1975, 1977). However, critics (Brodie III, 1993; Brodie III & Brodie Jr., 2004) of the clay replica studies have long argued that one needs only to encounter a single coral snake in nature to realise that motionless replicas are not testing a critical component of coral snake anti-predator defence. When a coral snake is threatened, it rarely flees directly: instead, it produces a vivid *in situ* thrashing display with kinked necks and curled tails that simultaneously makes it difficult to track the head and draws attention to the elevated tail, which in some species is combined with auditory cloacal “popping” (Moore, Danforth, Larson, & Davis Rabosky, 2019; Roze, 1996).

Although components of this innate behavioural display have been qualitatively reported in a number of colubrid snakes with mimetic coloration (Brodie III & Brodie Jr., 2004; Greene, 1979, 1988; Martins, 1996; Martins & Oliveira, 1998), no studies have systematically quantified variation in snake anti-predator displays across evolutionary origins or losses of mimicry. As a consequence, the use frequency and co-occurrence of “coral snake-like” display components are unknown. For example, do kinked necks always co-occur with curled tails, or do snakes often use only one of these components at a time? How do these correlation coefficients vary within and among species of models and mimics? Do closely related non-mimics ever use these display components, too? The impact of answering these and related questions is the ability to quantitatively measure the extent of mimicry between species and compare behavioural similarity across independent origins of any colour phenotype. Analysing behaviour within a phylogenetic comparative methods framework offers an exceptional opportunity to model the evolution of innate behavioural displays and statistically test hypotheses about how and why multi-component signals vary across species.

### Quantitative challenges of behavioural analysis in snakes and the comparative approach

Historically, there have been significant challenges to the quantitative and comparative study of snake non-locomotory behaviour, which often lacks analogy to existing methods designed for other taxa or purposes. From a biomechanical perspective, most motion analyses are based on locomotion, which involves translocation of a body that a) rarely obscures itself and b) demonstrates predictable, repetitive movement (Hu, Nirody, Scott, & Shelley, 2009; Jayne, 1988; Moon & Gans, 1998) These assumptions are rarely met for non-locomotory behavioural displays in snakes (*e.g.*, tight balling with a hidden head, death feigning). Automated visual tracking of full-body movement, as in studies of *Drosophila sp*. (Dankert, Wang, Hoopfer, Anderson, & Perona, 2009), is particularly difficult because snakes present as curved splines subject to nearly unconstrained deformations (*i.e.*, snake bodies can compress or inflate in all directions) and self-occlusions resulting from extreme body curvature. While affixing passive or active markers can aid in collecting motion data without these assumptions, the structured lipid layers throughout snake skin (Torri et al., 2014) inhibit many adhesives for temporary markers, and a permanent implant approach damages individuals that may be released or vouchered as museum specimens. These challenges have restricted data collection and analysis, so classical behavioural ethograms or presence/absence scores have been favoured for studying snake behaviour (Arnold & Bennett, 1984; Ford, 1995; Greene, 1979; Herzog & Schwartz, 1990).

However, macroevolutionary comparisons across clades require a modified approach. Unlike mammals or birds, in which entire orders differ primarily in the frequency and order in which they perform a shared set of behaviours (Reznikova et al., 2019; Winger, Barker, & Ree, 2014), even closely related snake species can differ greatly in the behaviours used (see Supplementary Material Video 1 for an overview of this diversity). This high behavioural diversity with little to no overlap in character states among species renders highly tailored ethograms broadly inapplicable across large clades. Additionally, many scoring systems use subjective descriptions of behavioural states that result in high user bias, such that two observers independently scoring the same trial (or its video recording) may create very different ethogram profiles (Burghardt et al., 2012; Donat, 1991). This user effect is driven primarily by the level of detail scorers provide, as some viewers make ethograms very complex with many transitions among detailed states, resulting in the majority of statistical variation being apportioned to differences in score complexity rather than content. Ideally, the best analytical framework for systems with these challenges should have low variance across individuals scoring the same trial and be infinitely expandable to account for novel behaviours without losing ability to compare to known behaviours (Ford, 1995; Podsakoff, MacKenzie, Lee, & Podsakoff, 2003).

Here we propose a new method for scoring snake anti-predator displays with both low observer bias and infinite extension, and then we show its utility in comparing the behaviour of highly divergent Neotropical snake species. First, we detail our approach for creating highly portable, pop-up arenas that standardise field collection of video data for studying snake behaviour or biomechanical kinematics (Figure 1). Next, we present a characterisation of anti-predator displays across 20 snake species from the Peruvian Amazon using our scoring model to demonstrate its implementation and analysis. Finally, we apply these new behavioural data to the study of anti-predator signal evolution within a phylogenetic comparative methods framework.

**Figure 1.**
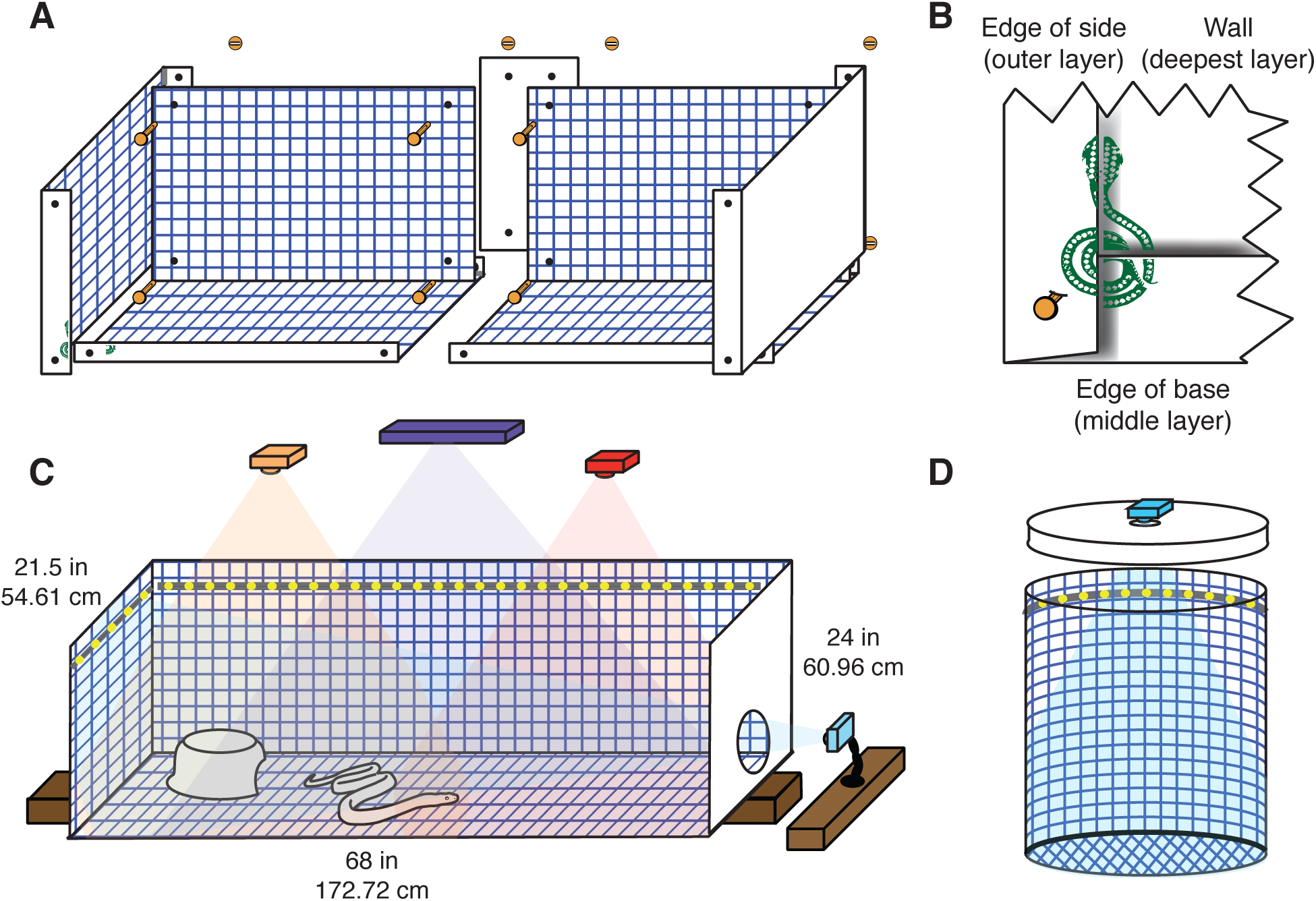
Diagrams of arenas used for measuring anti-predator response in the field. **A.** 10 pieces of corrugated plastic were connected using 16 brass fasteners and washers (the closest side walls and support piece are not shown, see Supplementary Figure S1). **B.** Close-up of lower left corner of arena showing detailed assembly. Each corner was labeled with an asymmetrical design to aid in proper assembly. **C.** GoPro cameras and respective fields of view are shown in orange, red, and blue within the assembled arena. Kinect camera (data not used in this study) is shown in purple. Cameras should be mounted on tripods or wooden planks that are independent of the arena to minimise camera movement. The wooden plank beneath the arena can be used to transmit vibrational stimuli. **D.** Buckets are labeled with gridlines to collect video data during field encounters. A hole is cut in the lid of the bucket and clear plastic covers the hole to enable videography while minimising the risk of escape or interference from rain. A GoPro camera (blue) is mounted on the lid of the bucket. Strings of LED lights are represented by yellow dotted grey lines around the tops of the arenas in panels C and D.

In combination, we hope that the convenience and flexibility of our standardised methodology and analytical model inspire researchers to 1) proactively collect more behavioural data on snakes in the field and to 2) expand our model for macroevolutionary analyses across more organisms with otherwise intractable behavioural diversity.

## Methods

### Field collection

We measured the anti-predator response of field-captured snakes (Table 1) across four field stations in the Amazonian lowlands of eastern Peru between March 2016 and December 2018: Villa Carmen Biological Station (850m elevation, latitude: -12.89, longitude: -71.40), Los Amigos Biological Station (270m elevation, 12.56, -70.10), Santa Cruz Biological Station (100m elevation, -3.52, -73.18), and Madre Selva Biological Station (100m elevation, -3.69, -72.46). We collected snakes through a combination of standardised drift-fence lines with both pitfall and funnel traps (Rabosky et al., 2011) and hand-foraging during both the day and night. All field methods were approved by the University of Michigan Institutional Animal Care and Use Committee (Protocols #PRO00006234 and #PRO00008306) and the Servicio Nacional Forestal y de Fauna Silvestre (SERFOR) in Peru (permit numbers: 029-2016-SERFOR-DGGSPFFS, 405-2016-SERFOR-DGGSPFFS, 116-2017-SERFOR-DGGSPFFS).

**Table 1.** Individuals of 20 Neotropical snake species collected in Peru. Abbreviations: Museo de Historia Natural de la Universidad Nacional Mayor de San Marcos (MUSM), University of Michigan Museum of Zoology (UMMZ), Male (M), Female (F), Juvenile (J), Villa Carmen (VC), Los Amigos (LA), Santa Cruz (SC), and Madre Selva (MS) Biological Stations.

| Family | Species | Field Number | Catalog Number | SVL (mm) | Mass (g) | Sex | Temp. (°C) | Locality |
| --- | --- | --- | --- | --- | --- | --- | --- | --- |
| Typhlopidae | *Typhlops reticulatus* | RAB2918 | UMMZ-248453 | 313 | 16.97 | F | 25.6 | LA |
| Aniliidae | *Anilius scytale* | RAB1140 | MUSM-36848 | 495 | 14.27 | F | 24.6 | LA |
| Boidae | *Corallus hortulanus* | RAB3499 | UMMZ-248448 | 1359 | 490 | F | 26.9 | LA |
|  | *Epicrates cenchria* | RAB2268 | not vouchered | 1050 | 1350 | F | 25.1 | SC |
| Viperidae | *Bothrops atrox* | RAB589 | MUSM-36921 | 483 | 37 | J | 32.0 | VC |
|  | *Lachesis muta* | RAB3562 | UMMZ-248454 | 1000 | 375 | F | 27.7 | LA |
| Elapidae | *Micrurus annellatus* | RAB3275 | UMMZ-248451 | 497 | 18.11 | F | 25.0 | LA |
|  | *Micrurus lemniscatus* | RAB3487 | UMMZ-248452 | 632 | 28.44 | F | 24.7 | LA |
| Dipsadinae | *Atractus elaps* | RAB3405 | MUSM-39784 | 367 | 14.67 | M | 25.0 | LA |
| (Colubridae) | *Dipsas catesbyi* | RAB3482 | MUSM-39823 | 435 | 19.64 | F | 25.6 | LA |
|  | *Helicops leopardinus* | RAB1811 | MUSM-37159 | 507 | 110 | F | 28.4 | MS |
|  | *Imantodes cenchoa* | RAB602 | MUSM-37248 | 370 | 4.17 | J | 28.9 | VC |
|  | *Ninia hudsoni* | RAB648 | UMMZ-248364 | 341 | 13.4 | F | 26.0 | VC |
|  | *Oxyrhopus melanogenys* | RAB3557 | UMMZ-248449 | 474 | 19.1 | M | 25.8 | LA |
|  | *Oxyrhopus petolarius* | RAB3406 | UMMZ-248450 | 338 | 9.37 | M | 25.4 | LA |
|  | *Thamnodynastes pallidus* | RAB1672 | UMMZ-246848 | 442 | 15.85 | M | 26.7 | MS |
|  | *Xenodon severus* | RAB608 | UMMZ-248367 | 511 | 90 | F | 28.4 | VC |
| Colubrinae | *Chironius fuscus* | RAB2669 | MUSM-38942 | 941 | 200 | M | 26.4 | LA |
| (Colubridae) | *Drymoluber dichrous* | RAB583 | MUSM-37089 | 391 | 19.72 | J | 27.0 | VC |
|  | *Leptophis ahaetulla* | RAB1670 | UMMZ-246824 | 684 | 53 | F | 27.1 | MS |

### Recording snake behaviour during capture

To ensure that initial anti-predator response was recorded at moment of capture, field survey personnel carried 20L buckets (38.50 cm tall, 34 cm diameter) during all transect or trail surveys and trapline checks. We designed two versions of these “bucket arenas” (Figure 1D): a lidded version for venomous snakes with a GoPro Hero 4 or 5 camera in a waterproof case mounted facing down over a hole cut in the lid that was covered with clear packing tape, and an unlidded version in which we simply hand-held the GoPro over the top of the bucket. For surveys in low light conditions, we lit the bucket’s interior either by shining 1-2 headlamps into the centre of the bucket floor or by lining the internal rim below the lid with a self-adhesive string of LED lights run by a custom-soldered waterproof battery pack (Lighting EVER, 5mm x 5mm Surface Mounted Diodes, 12V, 720lm/m). As soon as each snake was safely captured or removed from a trap, we placed it into the bucket and recorded behaviour for up to one minute at 120 frames per second (fps) and with a resolution of at least 1080 pixels. After filming, we then safely secured the snake within a cloth bag (and additional lidded bucket if venomous) for transport back to the field laboratory.

### Behavioural experimentation in the field laboratory

To balance our ability to capture the most biologically natural displays in a field setting with the experimental manipulations possible within laboratory infrastructure, we paired the bucket arena trials above with an additional set of trials in a “runway arena” set up at each field station. The constructed arena (Figure 1A-C) was made of 6mm thick white corrugated plastic (Corrugated Plastics, Hillsborough, New Jersey, USA) with the following dimensions: 1.73m long, 39.6 cm wide, and 61cm tall (see Figure S1 for individual piece sizes and overlap folding). This arena is made of 10 assembled pieces, so that it can be broken down and carried in a gear bag meeting standard airline luggage specifications (24” x 36” artist portfolio bag if desired to carry it separately from other field gear). Pieces are joined in the field using 2” brass fasteners and washers such that the base folds over the long side panels, and the ends wrap around the outside of the base and side panels (Figure 1B). Attachment of the two halves and additional rigidity along the centre of the long side panels is provided by support panels attached at the centre seam. To aid both movement tracking within the arena and the correction of camera lens distortion and parallax, we marked each internal surface with a standard fiducial calibration grid. Although we have used several different scales and colours of these markings, we have found that the most convenient colour and spacing combination for downstream analysis is to use ≈2mm thick lines that are blue (a rare colour for snakes, and therefore easiest to remove via automated analysis if needed) at a 2-3cm square grid size (commercially available self-adhesive contact paper with machine-printed square blue grid lines is the simplest option). Upon assembly in the field station laboratory, we lined all internal joints with silicone to ensure that small snakes could not become trapped in any joint gaps.

To record behaviour, we used a set of 2-3 GoPro Hero 4 or 5 cameras to film overhead views (located approximately 30cm from the short ends of the arena, 57 cm above the floor of the arena and directed downwards) and added a lateral view by mounting a camera flush against a transparent acrylic window in later models (Figure 1C). GoPro cameras can record synchronously using a Smart Remote (GoPro) accessory, or by synchronising all videos to an event that can be viewed in each camera field of view (*e.g.*, a ball bouncing or a laser pointer light). We provided consistent illumination across the arena by affixing a strip of flexible LED lights (same as bucket arena) to the top of the internal surface of the arena, which was powered either by a battery pack or by a transformer providing 12V of DC power from an AC electrical outlet. We also placed a Kinect V2 (Microsoft, Seattle, USA) perpendicular to the substrate to collect 2.5D depth maps of the snake behaviours (data not analysed for this study).

After capture, we kept snakes in cloth bags in a quiet, low disturbance part of the laboratory for between 12-24 hours and handled them as a little as possible between capture and completion of experiments in the field laboratory. We scheduled trials to reflect natural activity patterns by species, with diurnal snakes tested between 11am and 5pm and nocturnal snakes recorded from 5pm to 11pm. Each snake was then exposed to a standardised set of predator cues designed to simulate different information types snakes might use to assess predation risk. To simulate an avian attack from overhead, we created “looming” cues by swooping a black or white t-shirt above the arena to create both a shadow and pressure wave over the snake. A white shirt represented the countershading of a bird in the far distance, whereas trials with black shirts represented a bird passing over the animal at a close distance. To simulate the footfalls of an approaching large-bodied terrestrial predator like a mammal, we created “vibration” cues by playing a pulsed vibration (1 sec pulse, 1 sec no pulse) from a smartphone through a wooden plank underneath the bottom of the arena. To simulate a true attack by either predator class, we created a “tactile” cue in which we tapped the tail and body with a snake hook. While we realise that a warm body part may elicit different responses than a room-temperature metal object (Bowers, Bledsoe, & Burghardt, 1993), we used the snake hook to maintain consistency and safety across both venomous and harmless snakes. We randomised the order of cue presentation across individuals and recorded the snake’s response to each of these cues for one minute trials with one minute of rest in between trials. After the first trial in each set, we recorded the snake’s body surface temperature remotely without physical contact (which can change a snake’s behaviour and temperature, Schieffelin & de Queiroz, 1991) using an infrared temperature sensor (Raytek Ranger ST81). We cleaned the arena with unscented soap and water and dried it between each individual. We generally ran all arena trials with a team of two people, with one to administer the trials and one to timekeep, write notes, and operate the cameras. When testing any venomous snake, we required a third person whose sole job was to stand ready with a hook or tongs while continually monitoring and ensuring the snake’s safe containment.

All raw videos are permanently archived and publicly accessible on <u>deepblue.lib.umich.edu/data</u> [pending manuscript acceptance; for manuscript review purposes, please access them here: https://umich.box.com/s/v83zfb859s0nm9khr4zf132adikb9sa0].

### Behavioural scoring schematic

To score snake anti-predator displays from these videos, we designed a modular, objective schematic for assessing snakes’ behavioural states and their transitions. We identified our initial set of possible behavioural states by performing an inventory of snake behaviours observed in our dataset, conducting a literature review, and searching for videos of snake behaviours outside of our dataset on the internet. As one of our main questions involved correlated behaviours across body parts, we created separate sets of possible behavioural states for the head, the body, and the tail portions of the animal. The head and neck categories include distal portions of the body that may vary from frame to frame, as needed, to distinguish head and neck behaviours from body behaviours. Each ethogram is comprised of five separate categories: Shape, Presentation, Position, Posture, and Movement. Within each body part, we then scored each frame of each video across five categorical behavioural descriptors described below. Each category is infinitely expandable, but usually consists of three to eight non-overlapping states (see pictorial guides to observed states within the five categories; Figure 2 and Figures S2-S3).

**Figure 2.**
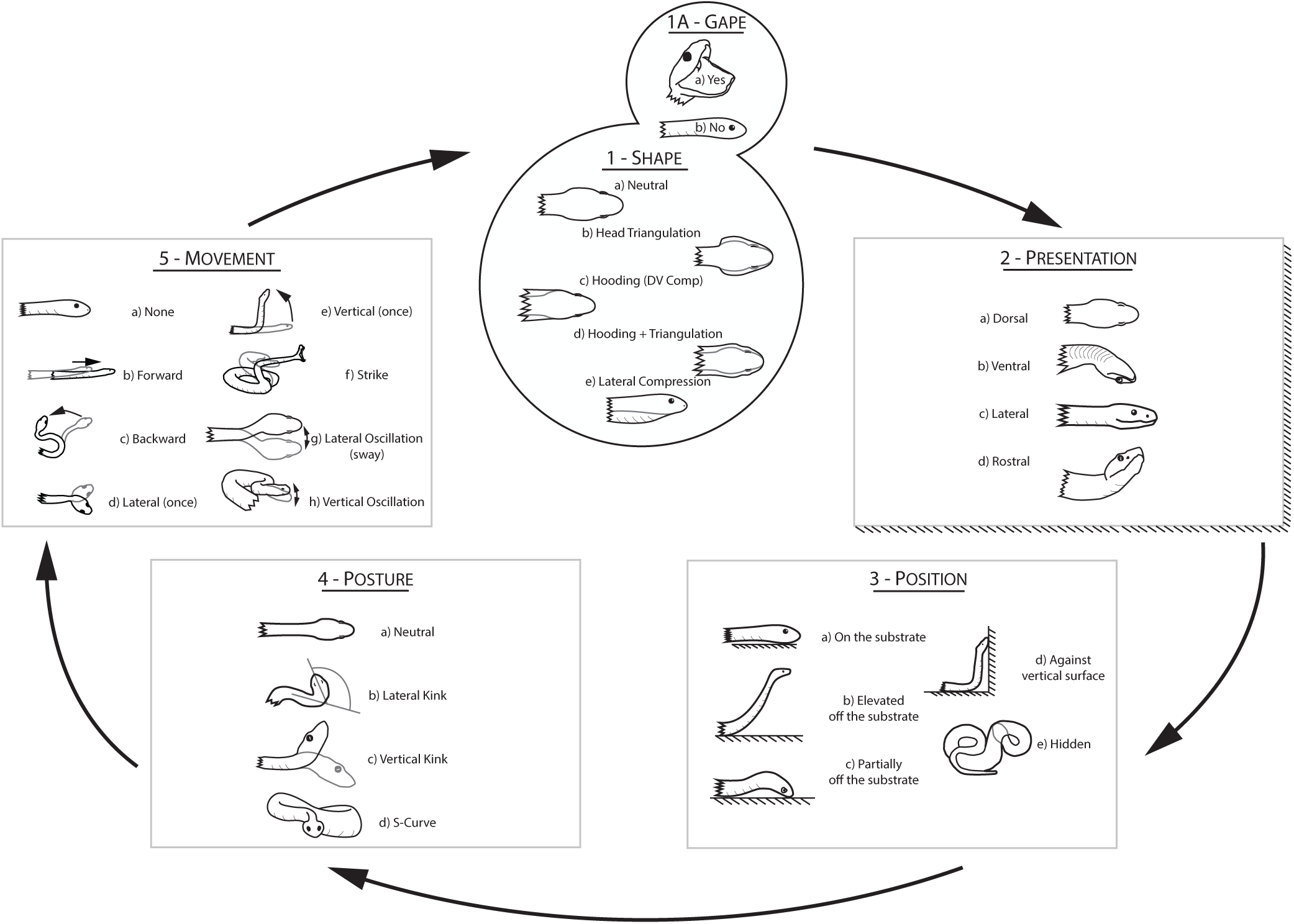
Modular schema for characterising behavioural states in snake heads. Our method objectively scores Shape, Presentation, Position, Posture, and Movement to test for state correlations among heads, bodies, and tails and which body parts are most responsible for convergence across taxa. Although states listed within each category are inclusive of known snake response variation, these categories expandable to accommodate future encounters with currently undescribed behaviours without losing similarity to the other categories. See Supplementary Figures S2-S3 for analogous schematics of body and tail behaviours.

We identified one state within each category for every frame of snake behaviour observed. Movement that was not obviously volitional (*e.g.*, falling) and frames in which a portion of the body were out of any camera view were not scored. A subset of example behaviours is provided as a supplementary video file (Supplementary Material Video 1).

1. **Shape** - describes how a body part deforms in dorsiventral and lateral dimensions from the normal resting posture with a cylindrical cross-sectional shape.
2. **Presentation** - describes which surface of the body part is facing upwards away from the substrate (which is consistent with the orientation towards the viewer in our trials or presumably towards a potential predator).
3. **Position** - describes the location of the body part with respect to the substrate.
4. **Posture** - describes the configuration of the body part in three dimensional space due to lateral or dorsiventral curvature, including kinking, coiling, or looping.
5. **Movement** - describes whether the body part is in motion and the manner in which it is moving, including directionality and repetition of movement if present.

To encourage consistency in behavioural scoring across individuals, all film observers scored a standardised “training set” of 10 videos before analysing data presented here. These multiply-scored training videos were used to test for the relative contribution of observer bias to the overall variance between scored ethograms (described below). Then, we scored one video for each of the 20 species analysed here to demonstrate examples of species-level variation elicited by our experimental protocol. Scorers watched all videos frame-by-frame using Quicktime Player 7 (Apple) and recorded observed behaviours in a Microsoft Excel template (Supplementary Material File 2) with drop-down menus to ensure standardisation to the common set of states described above (Figure 3). The majority of the behaviours were identified from a single camera view, although alternative views were consulted if the snake left the field of view or to identify behaviours such as dorsoventral flattening and partial elevation from the substrate.

**Figure 3.**
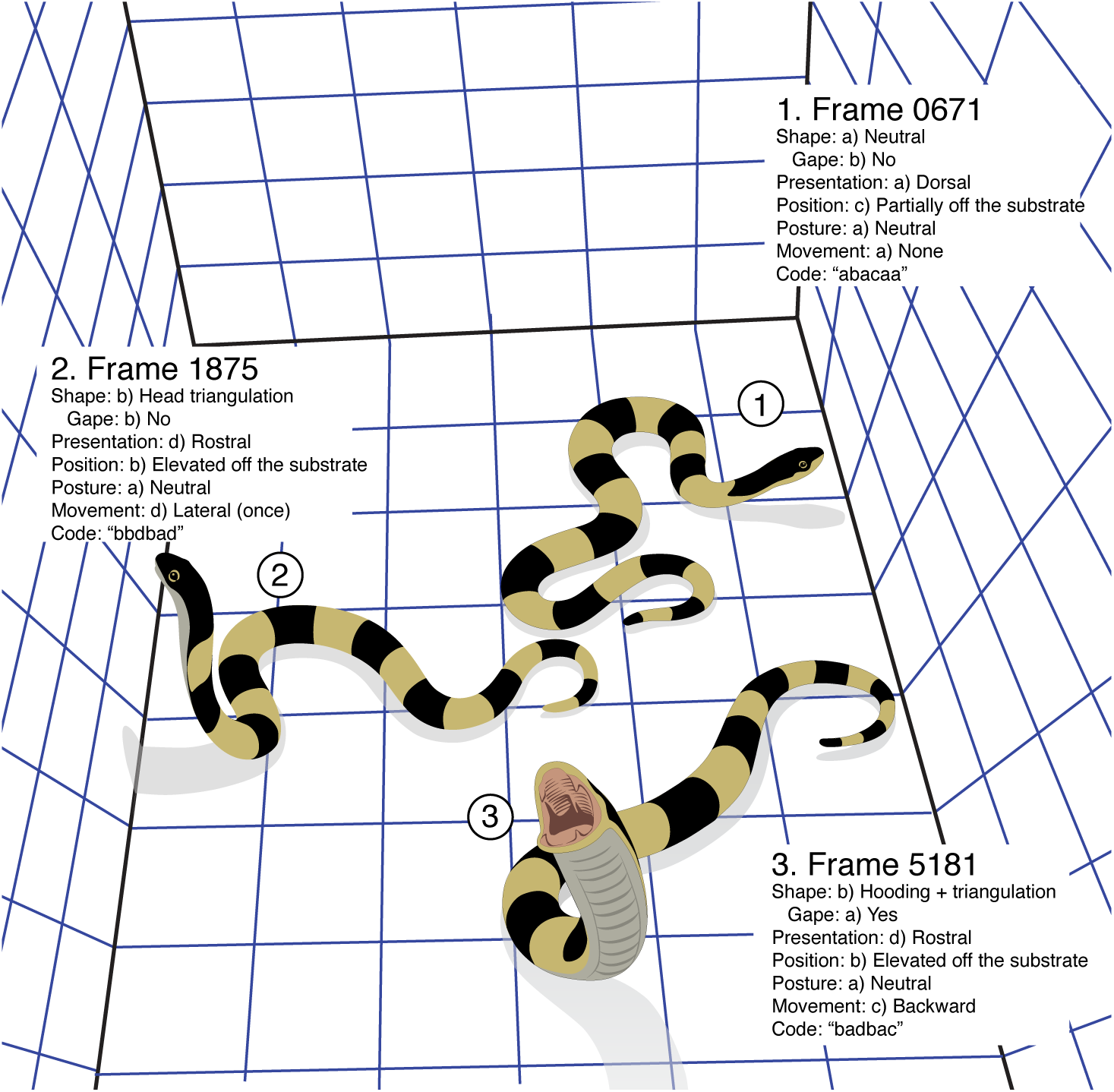
Illustration and scoring of three video frames from the exemplar trial for *Xenodon severus*, a non-venomous colubrid snake. White boxes show the state scores for the snake’s head at each frame, with all behavioural states corresponding to Figure 2. This snake employed hooding, gaping, and vigorous movement in response to the overhead looming stimulus in the later part of the trial. The full video of this trial can be found at <u>deepblue.lib.umich.edu/data</u>/pending (information pending manuscript acceptance; video for manuscript review at https://umich.box.com/s/efogjw6nrw1v3ncnc06b08o7cp0n73si).

### Statistical analysis

We performed all data handling and analysis in R v3.6.1, and all scripts are available with the scored ethograms on Dryad (information pending manuscript acceptance).

To compare across ethograms within the same body part, we first transcribed states for each body part into concatenated letter codes to create a vector with length equal to the total number of scored frames in that video (compare Figures 2 and 3 for an example of direct transposition of states to letter codes). We then calculated matrices of transition frequencies among body part states in each video frame and standardised them to allow comparison of videos of different lengths by dividing raw transition counts by the total number of frames in which the animal was visible (rather than by row sums as in a traditional frequency matrix). We chose to standardise frequency matrices in this manner to avoid over-inflating the frequency values of rarely observed states as a consequence of enforcing all row sums equal to 1, which was an undesirable statistical property in this context. In all other respects, these matrices were analogous to classical transition matrices.

To compare ethograms across videos that differed in the number and identity of states observed, we then transposed these values into a “global” matrix format that included the minimum set of all observed state combinations in this dataset. We chose this approach instead of one that included all possible state combinations because many states were never observed and would produce less computationally-tractable matrices (*e.g.*, 222 unique head state combinations observed in this dataset instead of 6400 possible head state sets). This transposition simply involved adding 0’s to state combinations observed globally but not locally for each ethogram. Finally, we computed pairwise distances between these matrices using three distance metrics (sum of the subtraction of one matrix from another, Euclidean distance, and Procrustes Similarity using the package “MatrixCorrelation” (Indahl, Næs, & Liland, 2018) and two weighting methods (weighted and unweighted) to compare metric performance. The weighting matrix had identical dimensions and states to the global transition frequency matrices, and each pairwise cell comparison varied systematically from 1 to 7 depending on the number of states that differed between the two concatenated letter codes (no difference = 1, one difference = 2, etc.). When used, we multiplied each transposed frequency matrix by the weighting matrix before computing the distance between the two matrices.

To assess the relative contribution of user bias to the overall variance in these comparisons, we used the unweighted Procrustes (PSI) distances between the ethograms to compare a) the same video scored by multiple people to b) different videos all scored by a single person. We used all pairwise comparisons of each type to generate and compare distributions and their summary statistics (*e.g*., median similarity value). If individual scorer bias is low, we would expect the similarity values of the same video scored by different people to be significantly greater (preferably 1, which would mean they are identical) than different snake videos scored by the same person. These distances independently calculated for heads, bodies, and tails were then also concatenated into overall similarity values for some downstream comparisons as follows.

To assess correlations between body parts, we measured both correlational coefficients of all state combinations and conditional probabilities based on a given behaviour. For example, to test how the head states correlate with a tail coil, we queried the tail data for all frames in which coiling occurs and graphed the frequency of each state as a stacked bar plot for each category. Thus, differences in head and tail correlation among species was discerned by comparing the stacked bar plots of head states corresponding to the same tail behaviour. To compare behavioural similarity among species to their taxonomic relatedness, we first calculated a distance-based phylogeny from the behavioural data for comparison to the time-calibrated molecular phylogeny of snakes (Pyron & Burbrink, 2014). We used the concatenated (heads, bodies, and tails) pairwise distances calculated by the unweighted subtraction method above and the package ‘phangorn’ (Schliep, 2011) to infer a neighbour-joining tree. Then we pruned the molecular phylogeny down to the taxa in our dataset and calculated similarity between trees using the Robinson-Foulds and path-based tree distance metrics in ‘phangorn.’

## Results

We found that this simple experimental setup (Figure 1) was very successful in eliciting and recording a diverse set of behavioural responses in snakes (see video highlights in Supplementary Material Video 1). Across both arenas, we observed many of the expected anti-predator displays known from the literature: a) viper-like head triangulation, gaping, striking, and puffing, b) coral snake-like kinked necks, periodic thrashing, body flattening, and tail curling, and c) ‘other’ known behaviours like fleeing, lateral tail vibration, hooding, and immobile coiling or balling (full videos deposited at <u>deepblue.lib.umich.edu/data</u>). However, we also observed unexpected behaviours with unclear functions, such as non-locomotory body undulations and jumping. We also found some individuals did not respond defensively, such as the bushmaster viper (*Lachesis muta*) included here that did not strike even when presented with a tactile stimulus. Critically, we found that much of the complexity in anti-predator responses was not readily apparent when simply observing these trials upon initial data collection, but this richness was very clear when watching and scoring the high speed videos afterwards.

### Processing and scoring speed

Although the number of snakes caught per day varied greatly (0 to 21) during our trips, we found that one team of two people running arena trials in the field could collect all standardised data at a rate of 6 snakes per hour (8 minutes for trials, 2 minutes for transferring snakes and cleaning the arena). As most collection days in the Neotropics yield 6 or fewer snakes per day, our high-throughput approach meant that collecting these data generally required just one hour per day (most efficiently run right before dinner, beginning with diurnal snakes) with minimal impact on other collection or survey activities. Bucket arena trials took even less time, with less than 2 additional minutes per snake capture (1 minute to ready the camera and transfer the snake to the bucket, 1 minute to film). Similarly, we found that after training, a typical video of approximately 5000 frames could be scored in a median time of 15 minutes, although the range varied from 3 minutes to approximately 6 hours, depending on video length and behavioural complexity.

### Measures of scorer bias

We found that the same video scored independently by multiple observers had much higher (2-3x) similarity scores than different videos scored by the same observer (Figure 4). However, these within-video similarity values averaged 0.75, which was lower than expected, although we did conservatively choose the training set to include species with only intermediate levels of behavioural differentiation. Overall, heads and bodies had much higher information content than tails, and correspondingly had larger differences between intra-scorer and intra-video distributions (compare Figure 4A and B to C). When we examined which of the 5 scoring categories contributed most to the variation among observers independently analysing the same video, we found that observers had the greatest discordance in scoring Position (proportion of variance in head scores, as an example: 0.295), Posture (0.203), and surprisingly, Movement (0.367). Together, these three categories accounted for more than 85% of the variance among scorers, and these relative contributions to variance were similar across body parts. Of particular scoring difficulty for all body parts was whether the snake was touching the substrate or not (Position), which was best solved by simultaneously examining both an overhead and lateral camera view of the same trial. Presentation scores had the least inter-scorer variance (proportion of variance for heads: 0.018).

**Figure 4.**
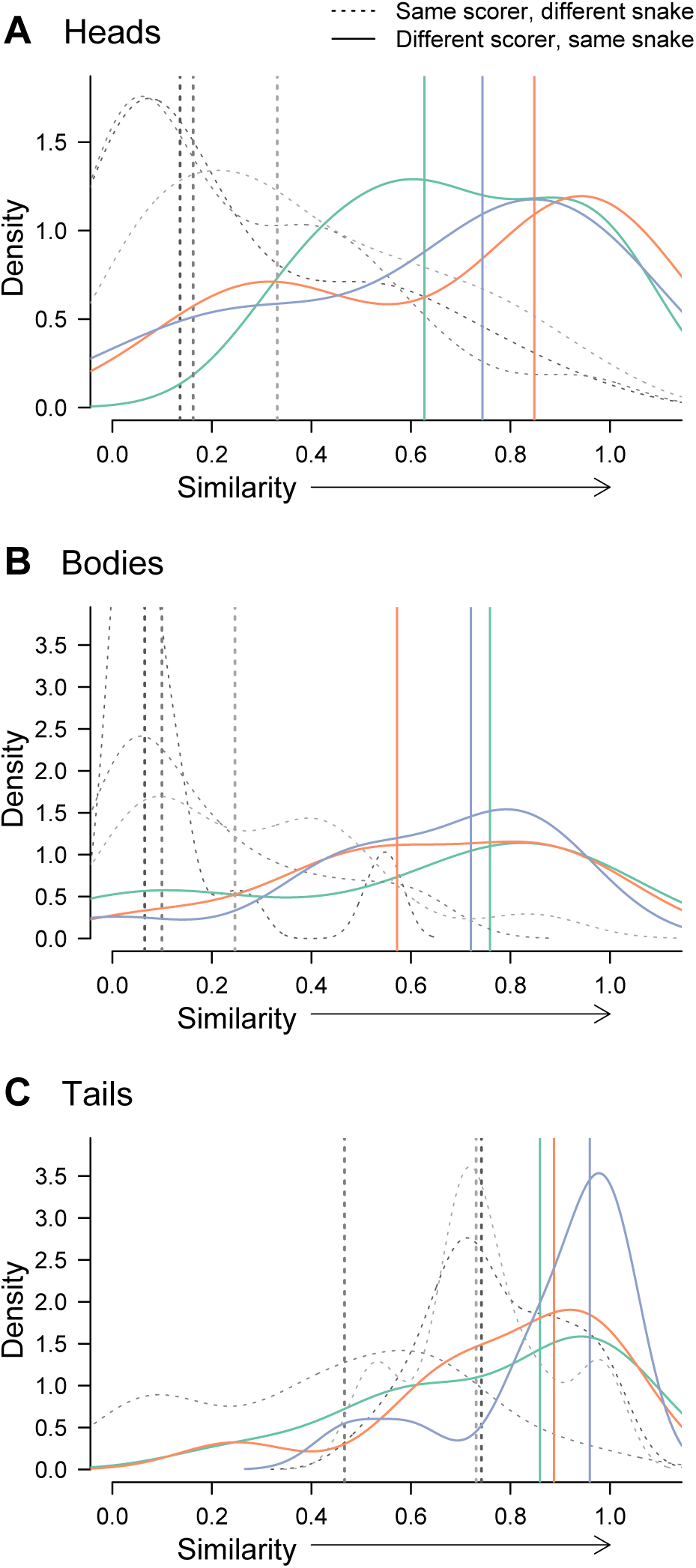
Estimates of scorer bias across videos. Density plots of pairwise ethogram comparisons across heads (**A**), bodies (**B**), and tails (**C**) demonstrate the relative contributions of scorers versus snakes to overall variance in ethogram scores. As expected, the same videos scored independently by different people (solid lines) are more similar than different videos scored by the same person (dotted lines). Vertical lines denote median values for each corresponding (by colour and line type) distribution of pairwise comparisons, and the comparison metric used is a Procrustes Similarity Index. Heads and bodies generally have higher information content and therefore greater discrimination capacity among species than tails.

### Correlations among body parts

We found that many state combinations that are theoretically possible within our scoring schematic were never observed, and the identity of unobserved states varied across species (Figure 5; all heatmaps archived at <u>deepblue.lib.umich.edu/data</u> [pending manuscript acceptance; zipped file for review at https://umich.box.com/s/vm4ccxpmg3yvvgh2vlb51cq39jb6kd3s]). In general, most individuals showed strong consistency in “stereotyping” their display across a trial, such that correlation coefficients across states were strongly bimodal (*e.g.*, each individual used a limited number of combinations, together, and consistently across a trial, yielding just a few combinations with high correlation coefficients; Figure 5). In particular, we found that the two coral snakes tested here (*Micrurus lemniscatus* and *M. annellatus*) produced vary similar displays to each other and that all presumed mimics tested displayed at least one component of the coral snake display (see archived heatmaps linked above). However, harmless snakes showed complex patterns of behavioural mimicry such that mimics could have just a tail display (*Anilius scytale*), just a neck kink (*Oxyrhopus petolarius*), a neck kink and thrashing (*Oxyrhopus melanogenys*), or thrashing and tail display (*Atractus elaps*). Both the extent to which the trial included these components and which were reliably used in combination varied greatly among mimics in comparison to non-mimics (Figure 5).

When complementarily evaluating the conditional probability of head states given a tail coil or kink, we again found that coral snakes and some mimics performed similar displays. Although tail coils were not exhibited by most species in our dataset, where possible these comparisons helped test which coral snake-like display components occurred more often than expected by chance in mimics versus non-mimics. For example, although the non-mimic *Chironius fuscus* displayed tail coiling and kinking, the head behaviours associated with these tail states differed greatly from coral snakes and colubrid mimics, especially in position, posture, and movement (Figure S4). This fine-scale analysis approach thus could both define which component states constitute the stereotyped display and discriminate among species that use these components differently, rather than simply testing for presence or absence of a feature. *Tests of evolutionary convergence*

**Figure 5.**
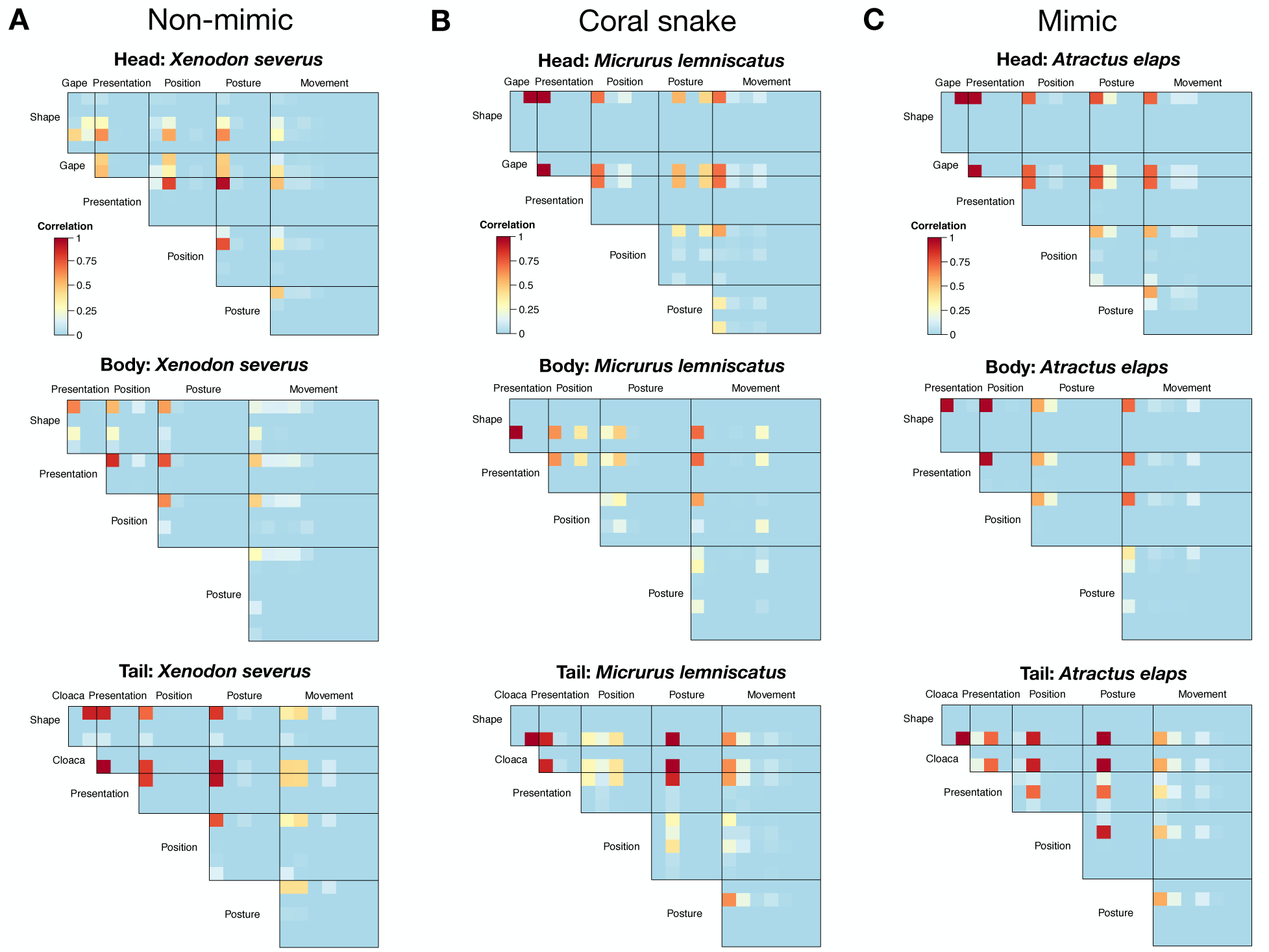
Correlation coefficients between states in models, mimics, and non-mimics. To visually represent a filmed trial in two dimensional print, heat maps of correlations demonstrate how much of the trial a snake spent in each combination of behavioural states across heads (upper panels), bodies (middle panels), and tails (lower panels). Warmer colours denote higher correlation coefficients, and the deepest blue colour represents state combinations never observed in the trial. Comparisons across a representative (**A**) non-mimic colubrid, (**B**) venomous elapid coral snake that serves as a model in a Batesian mimicry system, and (**C**) mimetic colubrid with black and red coloration that mimics a coral snake reveal not only strong behavioural similarity between the model and mimic, but also which behavioural components are responsible for that similarity and how often they are used during an anti-predator display. Heat maps for all species are archived with videos at <u>deepblue.lib.umich.edu/data</u> [pending manuscript acceptance; zipped file for review at https://umich.box.com/s/vm4ccxpmg3yvvgh2vlb51cq39jb6kd3s].

By evaluating behavioural similarity in comparison to the phylogenetic relationship of the species tested, we found clear evidence of both 1) behavioural convergence between coral snakes and their mimics and 2) closely related non-mimetic species that have particularly divergent responses, given their taxonomic similarity (Figure 6). Even though the mimics *Atractus elaps* and *Oxyrhopus melanogenys* represent different, evolutionarily independent origins of mimicry with each highly nested within clades of non-mimics (Figure 6, left), they have both converged on anti-predator displays that are more similar to *Micrurus* coral snakes than to any other snakes we tested. The individual of *Oxyrhopus petolarius*, in contrast, lacked strong behavioural similarity despite its red and black coloration. Additionally, the very distantly-related *Anilius scytale* (∼80 Ma divergence from Colubridae, Pyron & Burbrink, 2014) retains an anti-predator display that is more similar overall to other early-diverging lineages like blind snakes than to coral snakes, despite having both red and black coloration and some level of curled tail display. Conversely, several closely-related species of non-mimics displayed very dissimilar behavioural responses, including among the boas (*Epicrates cenchria* and *Corallus hortulanus*) and colubrine racers (*Leptophis ahaetulla* and *Drymoluber dichrous*; see also Supplementary Material Video 1). Lastly, we note that the different similarity metrics we used were variably responsive to different types of similarity, with some metrics producing behavioural trees of differing topologies (Robinson-Foulds distances varying from 30-32; Figure 6, right, shows simple unweighted pairwise matrix subtraction using all three body parts, RF=30). In addition, concatenated matrices of just heads and tails produced clearer evidence of convergence than those that also included bodies. Under all metrics, we recovered the behavioural convergence of at least one model and one mimic, as well as evidence of closely-related species dyads showing highly divergent behaviour.

**Figure 6.**
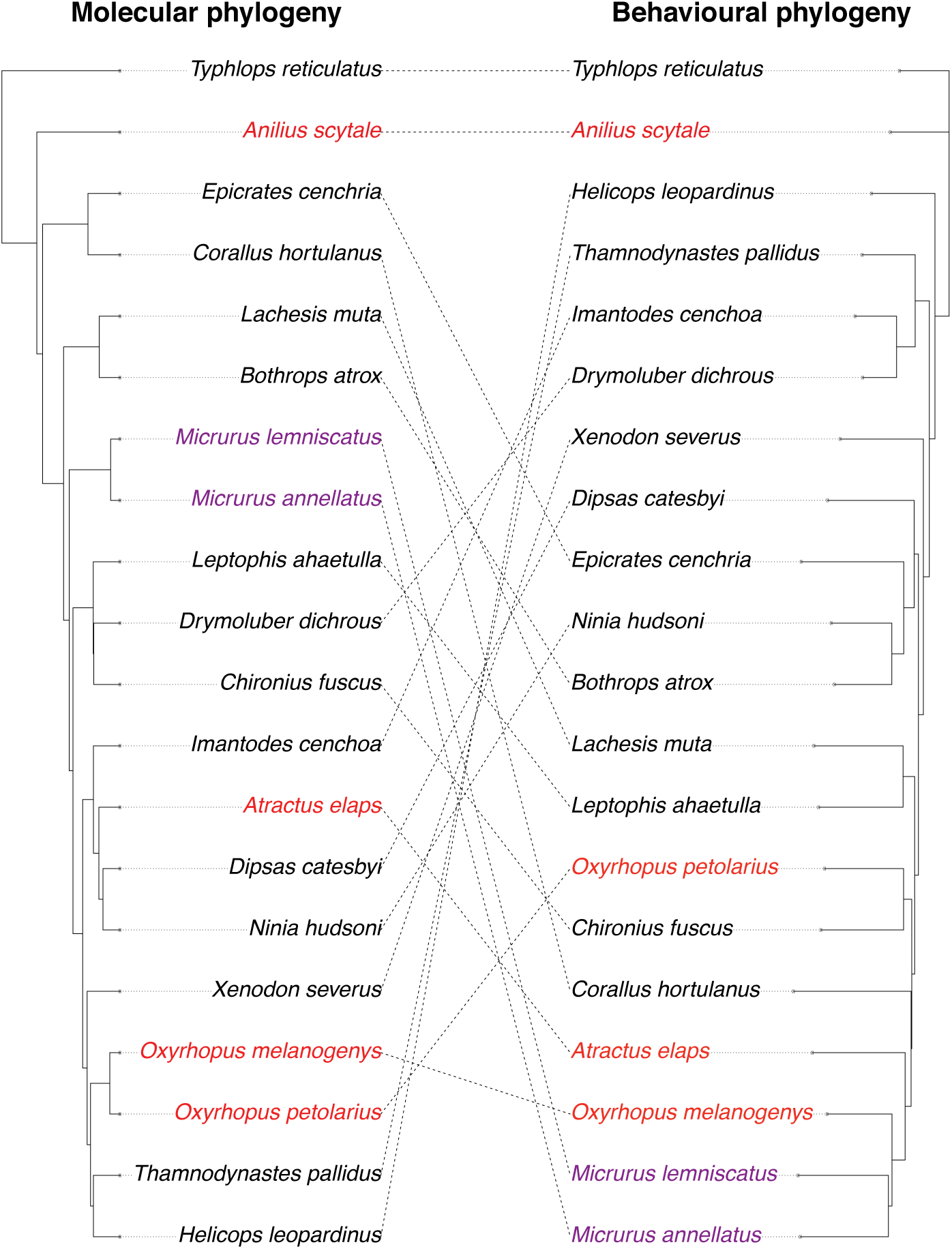
Evolutionary convergence in anti-predator displays between models (purple) and mimics (red). Co-plotted trees showing molecular (left) and behavioural (right) similarity of species demonstrates that across two independent origins of mimicry, colubrid mimics (*Atractus elaps* and *Oxyrhopus melanogenys*, in red) convergently evolved strikingly similar anti-predator displays to elapid models (*Micrurus lemniscatus* and *M. annellatus,* in purple). *Oxyrhopus petolarius*, in contrast, lacked strong similarity despite its red and black coloration. Additionally, the distantly-related *Anilius scytale* (∼80 Ma divergence from Colubridae, Pyron et al., 2013) retains an anti-predator display that is more similar overall to other early-diverging lineages like blind snakes than to coral snakes, despite having both red and black coloration and some level of curled tail display. Other closely-related species like the boas *Epicrates cenchria* and *Corallus hortulanus* employed very divergent anti-predator responses despite their taxonomic similarity. Behavioural phylogeny incorporates information from heads, bodies, and tails.

## Discussion

Here we have presented a new method for eliciting, quantifying, and comparing anti-predator displays across any species of snake collected in the field. We have also tested our method’s repeatability across independent observers and demonstrated some of the statistical analyses and questions addressable with data collected in this manner. In this exemplar set of 20 species from the Peruvian Amazon, we observed both classically-described and novel behaviours, as well as individuals performing displays in ways that were unexpected for their species given their phylogenetic placement. Given the high-throughput nature of our analytical approach and archived video data, these results have exciting implications for future analyses of snake behaviour, kinematics, and the evolution of anti-predator signals more generally.

### Performance of novel quantification approach and analysis

As we filmed snakes across five field expeditions spanning nearly three years, our approach to arena construction and experimental design underwent several rounds of revision. In our experience, the most important elements for a successful trial are a) testing snakes both at time of capture and across several experimental treatments, b) a well-lit arena to maximise video clarity, and c) a fiduciary grid with thin lines of a colour other than black so that the snake can always be discriminated from the background. Although we appreciated the greater space afforded by the runway arena because snake movement was less constrained and we could test larger-bodied species without escape, we also note that all four simulated predator cues can also be performed in the bucket arenas if the runway arena adds too much bulk or logistical complexity to a highly mobile field expedition. Most of the behaviours observed in the arenas were also seen in buckets, at least among the limited number of species analysed here. Overall, however, we were impressed with how well such a simple, portable, and high-throughput experimental design succeeded in eliciting and recording anti-predator displays across such a broad range of field conditions and snake species.

Our greatest analytical challenges were in developing a scoring method that could 1) accommodate high variation across body parts in any of the five components across species, 2) be infinitely expandable to include the previously undescribed, yet repeatedly observed, behaviours while retaining the ability to compare to similar behaviours, and 3) be implemented with low subjectivity across observers of any experience level. The ability to describe novel behaviours is particularly important when testing a species for the first time. With our method, novel behaviours can easily be described as new combinations of previously identified behavioural categories (*e.g.*, Position, Shape, Presentation) or by adding new states to categories as needed while retaining similarity across the other categories. By describing objectively observable physical states (*e.g*., which direction is the head moving, which surface is facing up) rather than subjective actions (*e.g.*, “exploring” or “death-feigning”), our ethograms limit subjective inference regarding the intent of the snake or whether the snake has entered or exited a particular “stereotyped display.” A single scorer’s assessment of this response may also change over time with experience (Dawkins, 2007). This subjectivity both within and among observers is mitigated by our system of objective descriptors that leave little capacity for alternate interpretations (Donat, 1991). Then, subjective decisions regarding which behavioural components must be present to qualify as a particular display can be made *ex post facto*, creating and analysing different composites based directly on scored data without needing to re-watch the video (Lehner, 1998). While our estimates of variance among scorers watching the same video was higher than we originally anticipated, the distribution of variance among categories (*e.g.*, higher in Position and lowest in Presentation) is valuable for informing how human perception influences interpretation of animal behaviour (Burghardt et al., 2012). We agree with these previous researchers (cited above) who state that quantitative assessments of user bias are underused but valuable analyses for behavioural studies.

Although our analytical methodology had many strengths, there are additional approaches that may be useful moving forward. Studies of complex, multi-component behavioural displays in other taxa often focus on modularity of correlated components (Rowe & Halpin, 2013; Scholes III, 2008). Other studies analyse complexity in sequence data using a Markov Process approach, especially if components are expected to follow a higher-order structure as in musical melodies (McAlpine, Miranda, & Hoggar, 1999) or other signals (Perry, Krakauer, McElreath, Harris, & Patricelli, 2019). These modalities may be analogous to communication via stereotyped behavioural repertoires so common among animals, and the application of similar Markov-based approaches to behavioural “phrases” in snakes may be an exciting future advance. Although there are significant challenges in quantitatively modelling behaviour in a comparative methods framework, they are similar to other multidimensional data types like geometric morphometrics (Zelditch, Swiderski, & Sheets, 2012) or diet (Grundler & Rabosky, 2019). Developing either a more modular or Markov approach might improve inference of behavioural trait evolution across the snake phylogeny, especially at a larger scale.

### Evolutionary convergence and divergence

We found clear quantitative evidence of behavioural convergence between coral snake models and some colubrid mimics, as well as examples of closely-related species showing surprising differences in anti-predator display. Although colubrid species with mimetic coloration are predicted to benefit from mimetic behavioural displays (Brodie III, 1993; Pfennig, Harper, Brumo, Harcombe, & Pfennig, 2007), this study is the first systematic quantification among species to assess how behavioural convergence varies across independent origins of mimicry. We have measured not only the behavioural similarity of models and mimics relative to other species (particularly closely related species with which they should share display components), but also assessed which behavioural components are driving that similarity. Our results suggest that the evidence for behavioural convergence between coral snakes and mimics is as compelling as the evidence for colour pattern convergence (Greene & McDiarmid, 1981), with similar levels of variation ranging from “somewhat similar” to “shockingly accurate”.

However, one important aspect of behaviour that we did not present in this study is an assessment of intraspecific variation (Bowers et al., 1993; Sealey, 2019). Although trait evolution studies often assign a single value to every tip/species in order to reconstruct rates of phenotypic change and a “species mean” approach is common for behavioural traits, intraspecific variation in anti-predator display absolutely exists and is critical to assessing character evolution (Omland, 1999). Although we did not analyse them here, we have observed a number of situations in which an individual’s response is highly context-dependent (*e.g.*, different displays elicited by different predator cues, Sealey, 2019) or in which some individuals readily perform a display while conspecifics do not (*e.g.*, some coral snakes never perform the thrashing display, Moore et al., 2019). The purpose of this study was to present a new analytical model and demonstrate its strengths and applications across an exemplar set of 20 species, but a more comprehensive future assessment of the evolution of anti-predator displays should both quantify and accommodate variation within and among individuals within single species.

### Implications for studying signal evolution

One of the most valuable applications of this work is in testing the relative contribution of colour pattern versus behavioural display in creating a successful signal that effectively deters a predator (Brodie III & Brodie Jr., 2004). How effective is a mimic that has an accurate colour pattern but low behavioural similarity in comparison to an inaccurate colour mimic with an accurate display? Are these two mimicry traits strictly additive, or does a holistic mimicry phenotype evolve and function in a way that is more complex than the sum of its parts? Previous work on the evolution of multimodal defensive signals (Rowe & Halpin, 2013) and their accuracy in mimicry systems (Dalziell & Welbergen, 2016) suggests that multimodality is so universal that aposematic coloration alone never creates a fully effective signal, but hypotheses about how and why these components co-evolve have not been broadly tested with empirical data. Using phylogenetic comparative approaches, and a much larger dataset, to reconstruct the timing, order, and trajectories of how colour and behaviour evolve across many independent origins of coral snake mimicry would create an explicit statistical framework for testing hypotheses about how and why both mimetic coloration and behaviour has evolved across species. This test would represent an exciting advance to our understanding of signal evolution within mimicry more generally, and coral snake mimicry is now the best candidate system for a tractable empirical test given statistical power derived from many identified independent origins within colubrids (Davis Rabosky et al., 2016).

In addition to advancing the quantitative analysis of snake anti-predator behaviours, this standardised method of video recording at the moment of capture enables detailed biomechanical examination of complex kinematic configurations that may not be observed in captivity. Such analysis can provide valuable data for studies such as constructing models of axial flexibility as a function of repeated vertebral elements (Galbusera & Bassani, 2019; Kambic, Biewener, & Pierce, 2017), understanding how non-legged vertebrates interface with complex substrates of varying surface properties (Abdel-Aal, 2013), and inspiring future generations of bio-inspired robotics (Danforth et al., in review; Marvi et al., 2014). This approach is key to rooting biomechanical and robotic studies more solidly in the context of ecology and evolution, while providing mechanistic explanations for the evolution of unique aspects of behaviour (Clark et al., 2019; Moore & Biewener, 2015). Additionally, these permanently-archived and publicly-accessible videos can be used for quantifying colour pattern, body size, sprint speed, and other aspects of basic snake biology. Given that knowledge about snakes is so highly dependent on low encounter frequencies, capturing any video data in semi-natural contexts represents an invaluable research contribution with many downstream applications and research opportunities. With current technology, quantifying behaviour across the entire snake tree of life is now limited only by the willingness of researchers to carry a gridded bucket and a GoPro while sampling in the field.

## Supporting information

Supplemental File 2 Ethogram Template

## Acknowledgements

We thank Dan Rabosky and Rudi von May for substantial assistance in organising and running the large-scale field expeditions necessary for specimen collection, as well as Conservación Amazónica - ACCA and Project Amazonas for logistical support at field stations. We also thank the many additional people who helped catch and film snakes in the field: Consuelo Alarcón Rodriguez, Arianna Basto, Amaranta Canazas Terán, Heidy Cárdenas, Yohamir Casanca Leon, Juan Carlos Cusi, Peter Cerda, Mark Cowan, Elar Durand Salazar, Maggie Grundler, Michael Grundler, Valia Herrera, Iris Holmes, Oscar Huacarpuma Aguilar, Edgar Iglesias Antonio, Eliz Lennia, César Macahuache Díaz, Daniel Nondorf, Greg Pandelis, Imani Russell, Roy Santa Cruz Farfán, Niery Tafur Olortegui, Tara Smiley, Pascal Title, Erick Vargas Laura, and Randi Villarcorta Díaz. We thank the undergraduate assistants who helped score videos: Sean Callahan, Ighodalo Eboigbe, Paul Hampel, O. Quinn Merchant, Aarcha Thadi, Elijah Thompson, Kayla Winter, and Emily Zuo. We also thank SERFOR (Servicio Nacional Forestal y de Fauna Silvestre) in Peru and US Fish and Wildlife service for scientific collection, export, and import permits and the MUSM in Lima for access to specimens. Finally, we thank John Megahan for scientific illustration of the trial shown in Figure 3. This research was supported with funding from the University of Michigan to ARDR and TYM and the Packard Foundation to Dan Rabosky.

