## Supplemental Figures S1-4 for "Convergence and divergence in anti-predator displays: A novel approach to quantitative behavioural comparison in snakes"

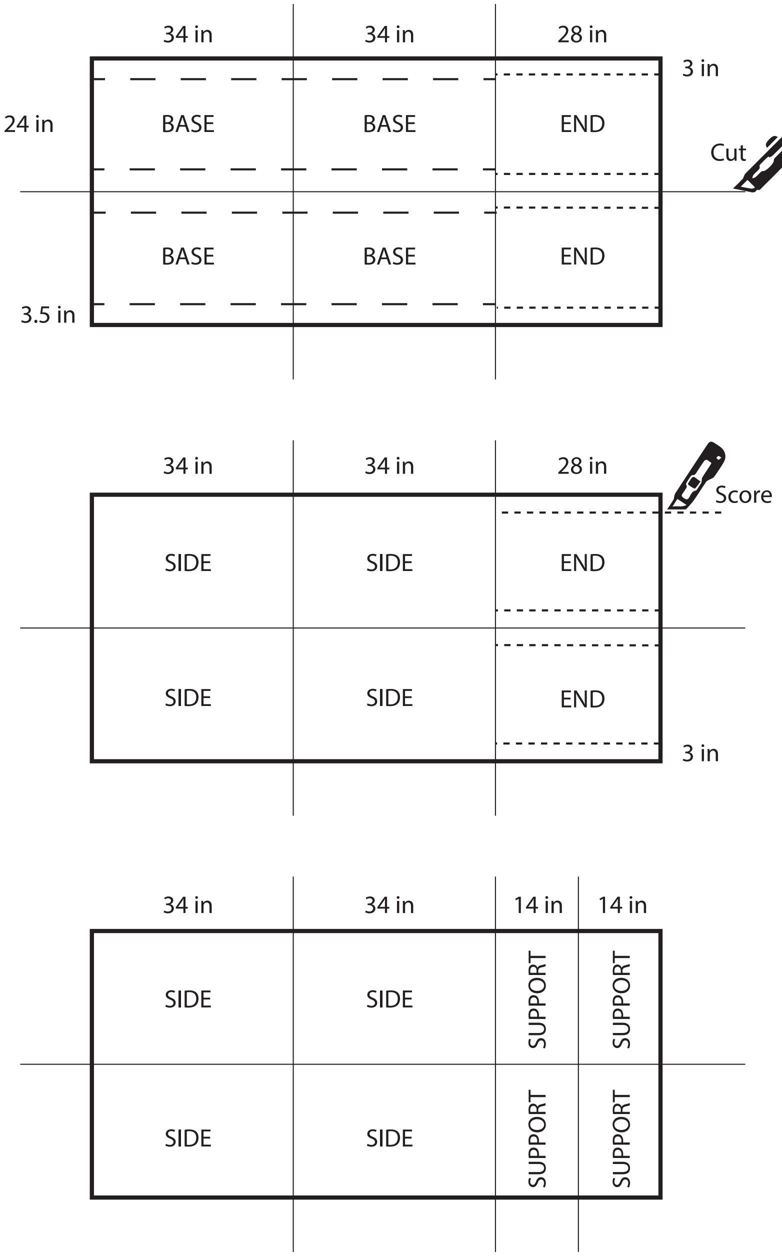


**Supplementary Material Figure S1.** The dimensions and the cutting patterns for each piece of corrugated plastic. This pattern produces two arenas from three pieces of 122cm x 244cm (48" x 96") corrugated plastic.

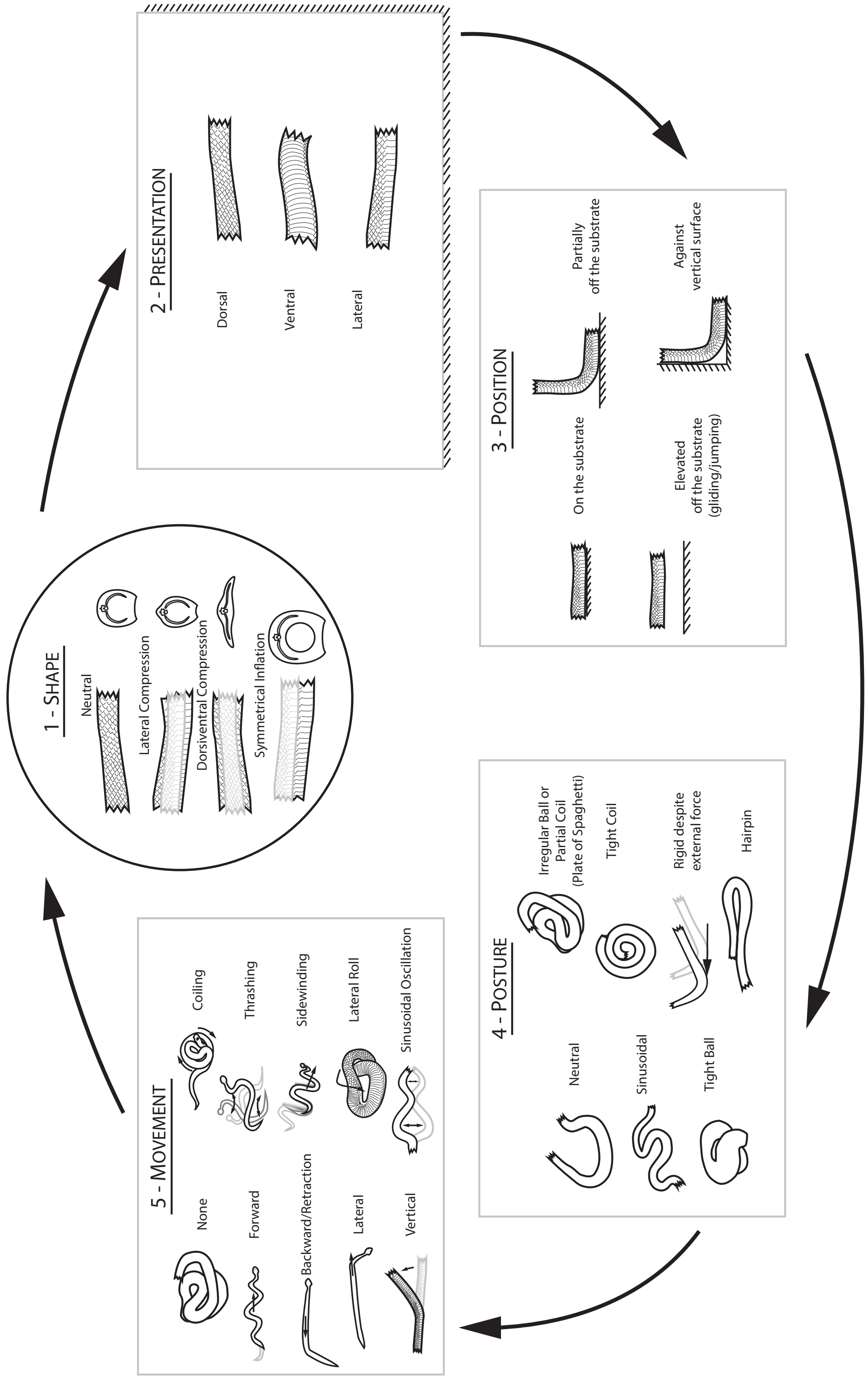


**Supplementary Material Figure S2.** Modular schema for characterising behavioural states in snake bodies by scoring Shape, Presentation, Position, Posture, and Movement for each video frame in a behavioural trial of anti-predator displays.

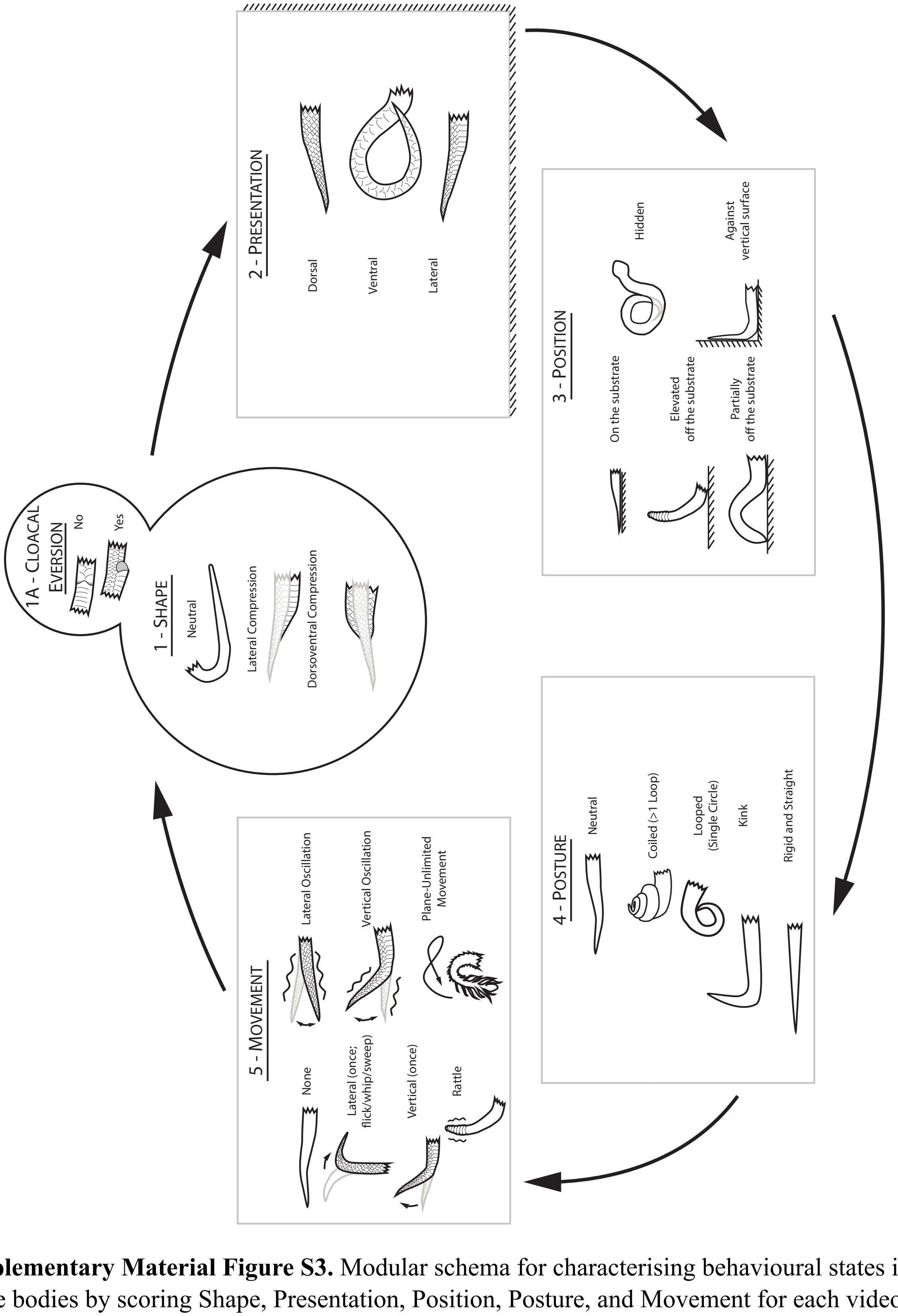


**Supplementary Material Figure S3.** Modular schema for characterising behavioural states in snake bodies by scoring Shape, Presentation, Position, Posture, and Movement for each video frame in a behavioural trial of anti-predator displays.

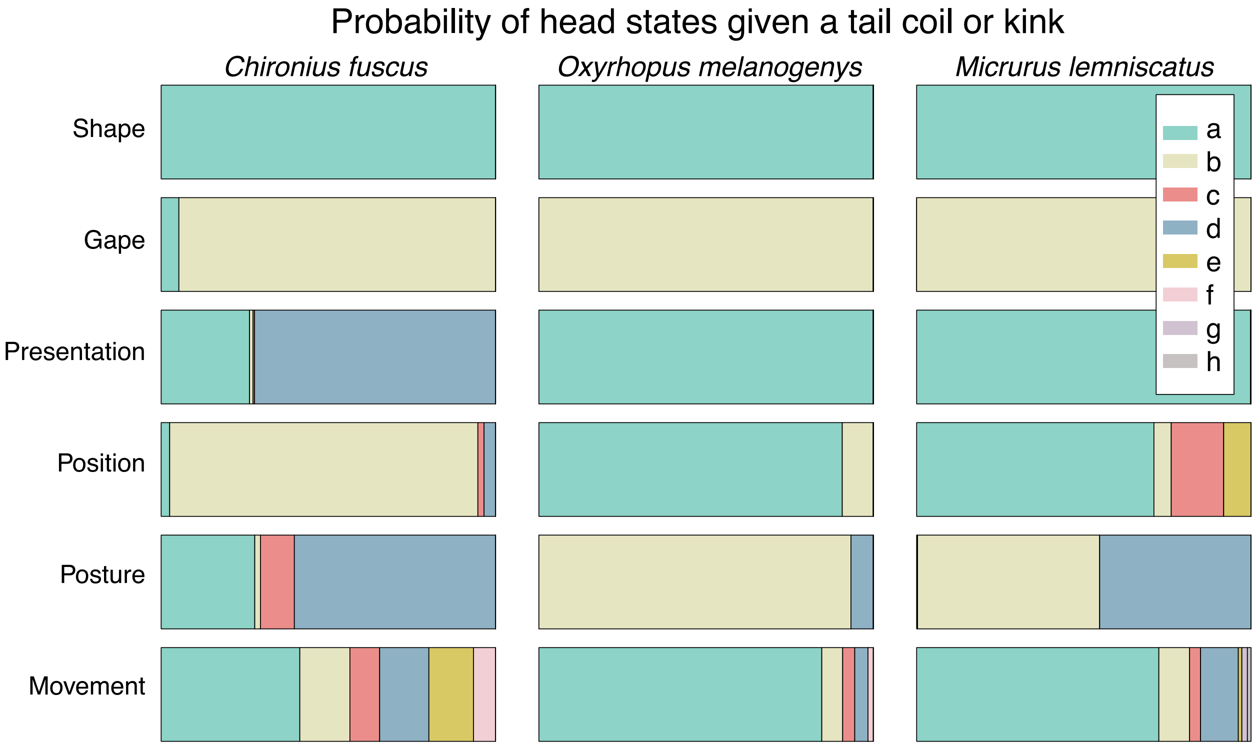


**Supplementary Material Figure S4.** The conditional probability of head states, given a tail coil or kink is shown as bar graphs for three species. Each behavioural category is represented by stacked bars representing the proportions of each state exhibited for that category. Tail coils were not exhibited by most species in our dataset. For the colubrid non-mimic *Chironius fuscus* (left), the head behaviours associated with tail coiling or kinking differed greatly from those of the elapid coral snake *Micrurus lemniscatus* and colubrid mimic *Oxyrhopus melanogenys*, especially in position, posture, and movement.
